# VNC-Dist: A machine learning-based semi-automated pipeline for quantification of neuronal positioning in the *C. elegans* ventral nerve cord

**DOI:** 10.1101/2024.11.16.623955

**Authors:** Saber Saharkhiz, Wesley Chan, Chloe B. Kirezi, Mearhyn Petite, Tony Roenspies, Theodore J. Perkins, Antonio Colavita

## Abstract

The *C. elegans* ventral nerve cord (VNC) provides a simple model for investigating the developmental mechanisms involved in neuronal positioning and organization. The VNC of newly hatched larvae contains a set of 22 motoneurons organized into three distinct classes (DD, DA, and DB) that show consistent positioning and arrangement. This organization arises from the action of multiple convergent genetic pathways, which are poorly understood. To better understand these pathways, accurate and efficient methods for quantifying motoneuron cell body positions within large microscopy datasets are required. Here, we present VNC-Dist (Ventral Nerve Cord Distances), a software toolkit that replaces manual measurements with a faster and more accurate computer-assisted approach, combining machine learning and other tools, to quantify neuron cell body positions in the VNC. The VNC-Dist pipeline integrates several components: manual neuron cell body localization using Fiji’s multipoint tool, deep learning-based worm segmentation with modified Segment Anything Model (SAM), accurate spline-based measurements of neuronal distances along the VNC, and built-in tools for statistical analysis and graphing. To demonstrate the robustness and versatility of VNC-Dist, we applied it to several genetic mutants known to disrupt neuronal positioning in the VNC. This toolbox will enable batch acquisition and analysis of large datasets across genotypes, thereby advancing investigations into the cellular and molecular mechanisms that govern VNC neuronal positioning and arrangement.

## 1. Introduction

Fluorescent reporters have become invaluable tools for investigating nervous system connectivity and function in *C. elegans* [1]. In the adult *C. elegans* hermaphrodite, the nervous system contains 302 neurons, organized into sensory neurons, interneurons, and motoneurons, with most cell bodies clustered in the nerve ring, retrovesicular ganglion (RVG), preanal ganglion (PAG), and along the ventral nerve cord (VNC) [2–4]. The position, morphology, and connectivity of different sets of these neurons can be readily visualized using distinct colours of cell-specific fluorescent markers [5]. However, accurately and reproducibly identifying and quantifying cells remains challenging and time-consuming, particularly when analyzing large sample sizes or conducting high-throughput genetic screens [6–8].

At hatching (L1 stage), the *C. elegans* VNC contains three classes of motoneurons: 6 GABAergic inhibitory DD, 9 cholinergic excitatory DA, and 7 cholinergic excitatory DB, arranged in a highly stereotyped manner that includes repeating DD-DB-DA motor pools innervating body wall muscles [9] (Fig. 1a, b). This organization is largely established between the bean and 2-fold stages of embryonic development, during which DD, DA, and DB progenitors, initially located on both the left and right, undergo convergent extension to move medially and position themselves along the anterior-posterior axis, while also engaging in poorly understood cell sorting, and separation movements that form the VNC [10]. Notably, aspects of this process share molecular and cellular mechanisms with neural tube formation in vertebrates, suggesting that the VNC may serve as a genetically tractable model for studying the underlying cell and molecular biology [11].

**Fig. 1.**
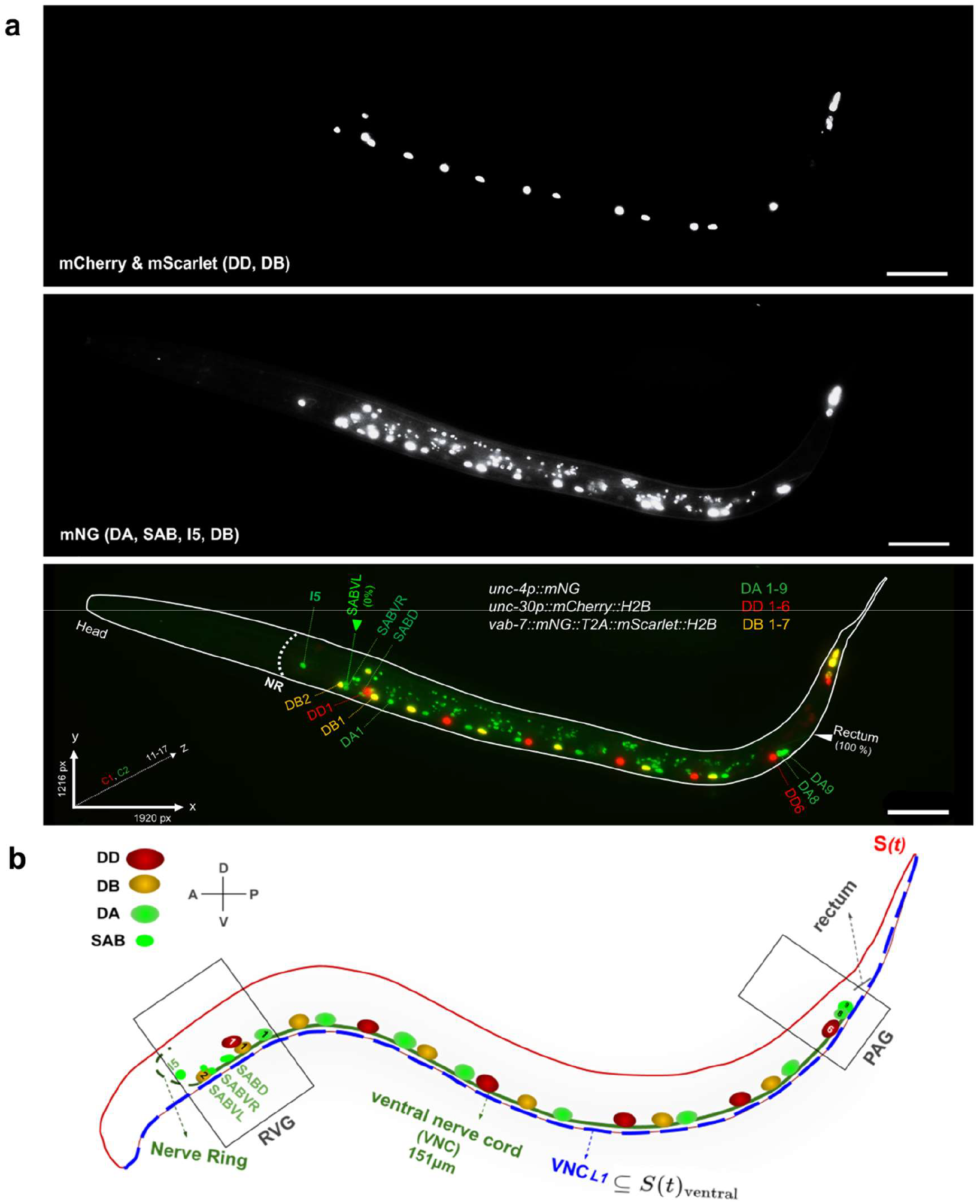
Ventral nerve cord of a newly hatched wild type *C. elegans* hermaphrodite. **(a)** Max-intensity-projected fluorescence micrograph of the *C. elegans* ventral nerve cord (VNC) expressing nuclear-targeted reporters: mCherry labels DD neurons (red), mNeonGreen labels DA, SAB, and I5 neurons (green), and co-expression of mScarlet and mNeonGreen labels DB neurons (orange). Tail epithelial cells also express the DB reporter. Reference points are SABVL (0 % anterior) and the rectum (100 % posterior); scale bar, 20 µm (1 µm= 6.82 pixels). **(b)** Diagram of a wild type (WT) *C. elegans* at L1 highlighting key structures: the nerve ring (NR), ventral nerve cord (VNC), DD, DA, and DB, SAB (SABVL, SABVR, SABD), and I5 neurons. The retrovesicular ganglion (RVG) contains SAB, DA1, DB1, DB2, and DD1 neurons; the preanal ganglion (PAG) contains DA8, DA9, and DD6. The worm’s body is modeled as a parametric spline function S(t) (red), with the VNCL1 spline (dashed blue) running along the ventral side of S(t), depicted in parallel to the VNC (green).

During embryonic VNC assembly, disruptions in the migration of DD, DA, and DB progenitors result in the misplacement of motoneuron cell bodies in newly hatched L1 larvae. One example involves disruption of a planar cell polarity-like pathway mediated by *vang-1*/VangL and *prkl-1*/Prickle, along with a parallel *sax-3*/Robo pathway, which together act redundantly to promote convergent extension of progenitors during VNC assembly [10]. Disruption of either pathway alone interferes with VNC assembly and causes motoneuron cell bodies to adopt abnormal positions at hatching. When both pathways are disrupted, convergent extension is severely compromised, resulting in a striking anterior displacement of most motoneuron cell bodies [10]. These findings indicate that motoneuron position and pattern in the L1 VNC provide a clear and accessible readout of defects that arise during embryonic VNC formation. Previous studies have primarily relied on manually measuring the positions of DD motoneurons [10]. However, a more complete understanding of VNC assembly requires accurately quantifying the position and pattern of all 22 DD, DA, and DB motoneurons [12–14]. To accomplish this, the DD, DA, and DB motoneuron classes must be distinctly labeled, and an efficient, high-throughput semi-automated workflow is needed to quantify their cell body positions along the VNC.

Current methods for quantifying complex curved biological structures, such as Neurite Tracer [15], NeuroQuantify [16], AniLength [17], SNT [18] and NeuronJ [19] typically depend on traced path analysis, generating low-precision discrete approximations that fail to capture continuous curve lengths. In contrast, Nellie’s pipeline employs hierarchical deconstruction and feature extraction to robustly analyze complex organelle architectures [20]. However, most of them demand substantial manual input and often fail to generalize to models like *C. elegans*, leaving researchers reliant on time-consuming manual tools, such as Fiji’s segmented line tool (Fig. 2) [21], which may lack consistency across users.

**Fig. 2.**
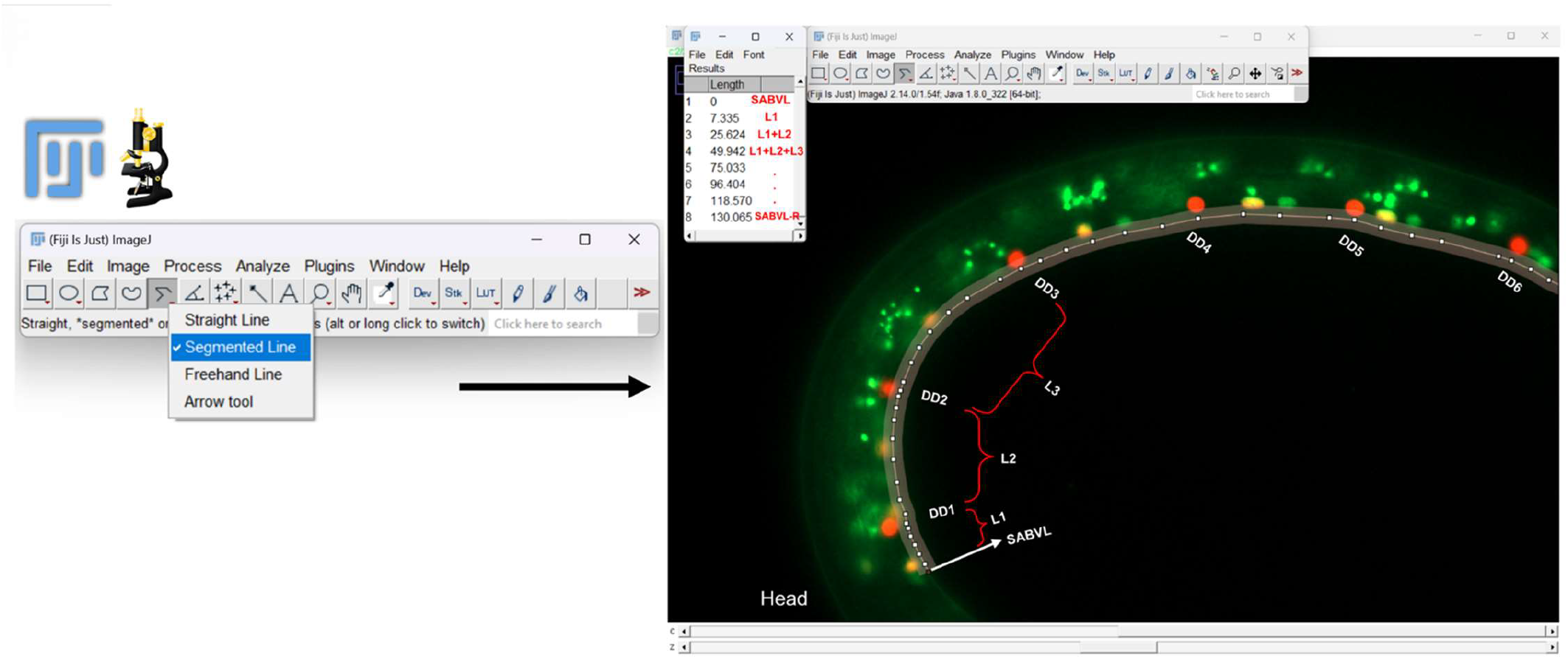
Established manual method for VNC motoneuron position quantification using Fiji’s Segmented Line tool. The analysis involves manually measuring distances through three separate traversals along the VNC for each motoneuron class: 3 SAB and 9 DA neurons (green), 6 DD neurons (red), and 7 DB neurons (orange).

Visually detecting shifts in VNC neuron positioning and patterning across genetic mutants or individual *C. elegans* is challenging due to the large number of neurons involved, the wide range of patterns that can result from disrupted VNC assembly, and the subtle or variably penetrant phenotypes that often accompany such disruptions [10]. Here, we introduce VNC-Dist (Ventral Nerve Cord Distances), a robust pipeline tailored for labs studying neuron positioning in *C. elegans*. VNC-Dist addresses these challenges through a mathematical algorithm that calculates the curve lengths along the *VNCL1* spline, which is a parametric curve tracing the VNC along the worm’s ventral outline [22](Fig. 1b). This pipeline enables rapid and precise batch analysis of 3D fluorescent images, significantly outperforming manual methods. We demonstrated its versatility across mutants like *vang-1/Van Gogh* and *sax-3/Robo*, which are known to cause VNC assembly defects [10]. VNC-Dist integrates SAM [23–25] for worm segmentation, OpenCV [26] for shape extraction and contour detection, and numerical analysis for accurate neuron-distance measurements [27]. We evaluated VNC-Dist with expert, intermediate, and beginner users and observed consistent learning curves in processing time, whereas manual length measurement was slower and highly variable. All users completed tasks more quickly and accurately with VNC-Dist, underscoring its robustness. This advantage stems primarily from the manual approach requiring numerous clicks per neuron class using Fiji’s Segmented Line tool [21] to trace the worm’s ventral edge, making it significantly more laborious for high-throughput analysis.

## 2. Materials and Methods

### 2.1 *C. elegans* strains, reporters, and imaging

#### 2.1.1 Strains

Worms were maintained at 20°C, and all strains were derived from the N2 genetic background. We used the following alleles and transgenes organized by the linkage group (LG): LGX: *vang-1(tm1422)* and *sax-3(zy5)*. LGII: *unc-4(zy123[unc-4::mNG])* and *zySi2[unc-30p::mcherry::H2B::unc-30]*. LGIII: *vab-7(zy137[vab-7::mNG::3xFlag])*, and *vab-7(zy142[vab-7::mNG::T2A::mScarlet-I::H2B])*. LGIV: *prkl-1(ok3182)*.

#### 2.1.2 Construction of DD, DA, DB fluorescent reporter strains

To label DD, DA, and DB motoneurons at the L1 stage, we used class-specific promoters to drive fluorescent reporters: *unc-30* (DD1–6), *unc-4* (DA1–9, SAB), and *vab-7* (DB1–7). *zySi2 [unc-30p::mCherry::H2B::unc-30 3’UTR + unc-119(+)]* was made using Mos1-mediated single-copy insertion (MosSCI) [28]. The MosSCI LGII ttTi5605 targeting and selection plasmid (pAC661) was constructed using Gibson Assembly to combine four PCR fragments: a ∼2.5 kb *unc-30* promoter from N2 genomic DNA, a ∼1.25 kb mCherry::H2B cassette from pOD2046 (Addgene #89367), a 410 bp *unc-30* 3’UTR from N2 genomic DNA, and a ∼7.4 kb MOS targeting and selection cassette from pCFJ151. CRISPR/Cas9 gene editing, utilizing the self-excising selection cassette (SEC) method, was used to knock-in mNeonGreen (mNG) at the C-terminus of the endogenous *unc-4* and *vab-7* genes [29]. Guide sequences for C-terminal insertion were identified using the IDT guide design tool at www.idtdna.com/site/order/designtool/index/CRISPR_SEQUENCE. The sequences were cloned into the Cas9–sgRNA plasmid pDD162 (Addgene #47549) using the NEB Q5 Site-Directed Mutagenesis Kit. *unc-4* and *vab-7* homology arms were cloned into the mNG-SEC plasmid pDD268 (Addgene #132523).

A co-CRISPR approach [30] was used to swap the 3xFlag sequence in *vab-7(zy137[vab-7::mNG::3xFlag])* with a *T2A::mScarlet-I::H2B* cassette to generate *vab-7(zy142[vab-7::mNG::T2A::mScarlet-I::H2B*zy137])*. A homologous repair plasmid (pAC798) containing 5’ and 3’ homology arms PCR amplified from *zy132* and a *T2A::mScarlet-I::H2B* cassette ordered as an IDT gBlocks gene fragment. An sgRNA targeting a site within the 3xFlag sequence was generated using pDD162 as described [29].

Primers are listed in Supplemental Table S1. Sequence files for pAC661 and pAC798 are included in the supplemental materials.

#### 2.1.3 Image acquisition

Images of VNC neurons were acquired using a Zeiss Axio Imager M2 widefield fluorescence microscope with a 40×/1.4 NA oil immersion objective. Images were saved as .czi files in a four-dimensional format (CZYX: two channels, 11–17 z-slices, 1216 × 1920 pixels). The channels and z-stacks were combined using maximum intensity projection and presented as 8-bit 2D images in TIFF format. Imaging resolutions were optimized for neuron diameters—mScarlet (0.258 µm), mNG (0.225 µm), mCherry (0.266 µm)—with an overall merged image resolution of approximately 0.266 µm and an isotropic voxel size of 0.146 μm. During image acquisition, worms exhibiting twisting, hooking, or coiling were excluded, as such deformations could compromise the segmentation accuracy and data reliability. For sample preparation, newly hatched L1 stage worms were mounted on 2% agarose pad slides containing 0.1 μg/μL of the paralytic Levamisole in M9 buffer.

### 2.2 VNC-Dist pipeline

#### 2.2.1 Overall pipeline design

Fiji’s multipoint tool is used to annotate neuron cell bodies and the rectum in anterior-to-posterior order using counter numbers (1: DAs, SABs, rectum; 2: DDs; 3: DBs), producing X–Y coordinates of all identified neurons and the rectum from the in-focus z-slices as a CSV file (Fig. 3a). The in-focus slices are then merged and collapsed into 2D TIFF images for segmentation by SAM-Plus. Finally, normalized relative distances along the ventral nerve cord’s anterior–posterior axis are computed by tracing the ventral boundary (*VNCL1*) (Fig. 1b).

**Fig. 3.**
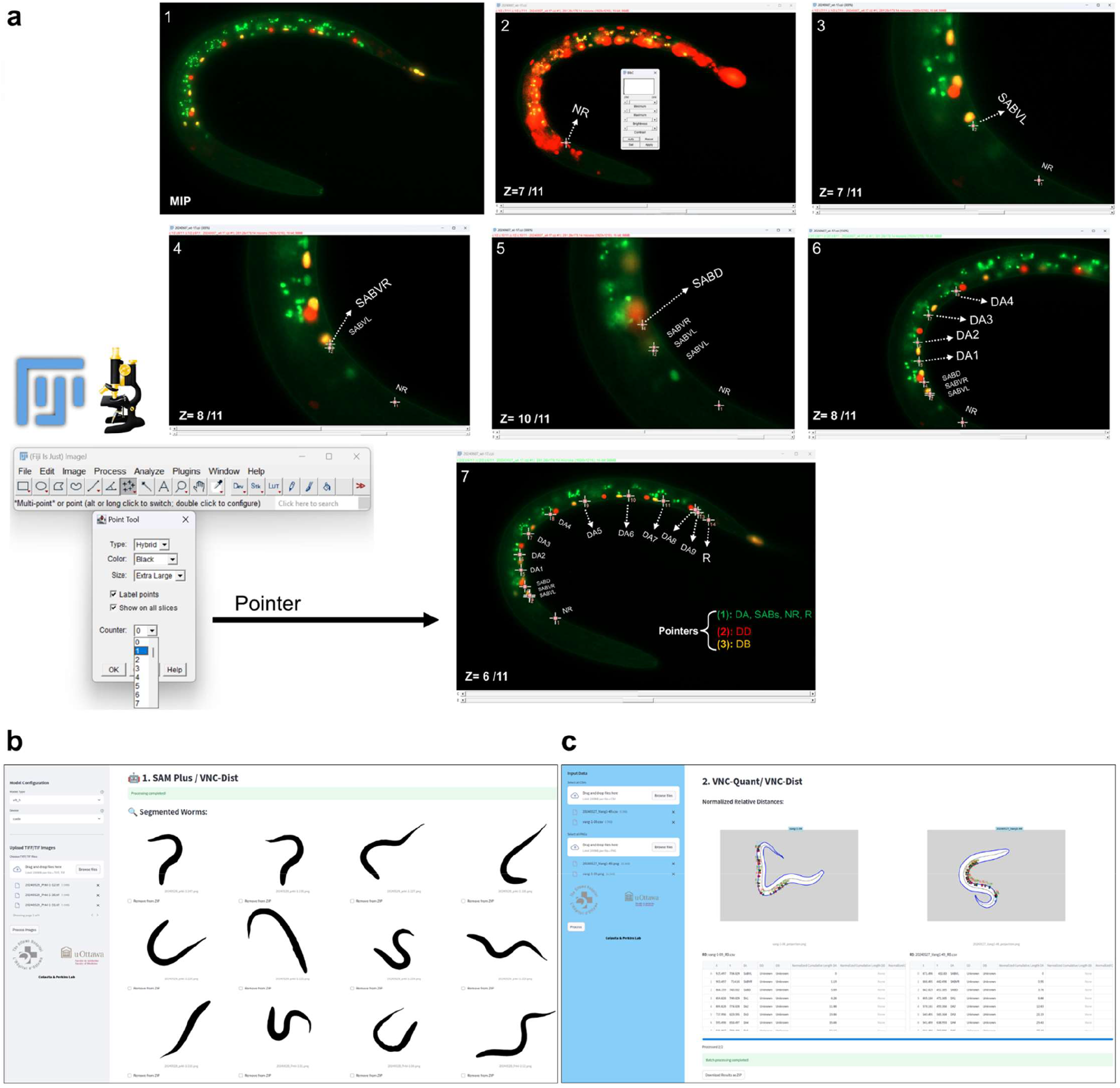
User interface of VNC-Dist pipeline for analyzing VNC neuron positioning. **(a)** Neurons between SABVL (0% reference point) to the rectum (100% reference point) are annotated using Fiji Annotator (multipoint tool) and classified into three groups based on Counter numbers (1: DAs, SABs, rectum; 2: DDs; 3: DBs). The X-Y coordinates are saved as a CSV file, and in-focus z-slices are merged and collapsed into 2D TIFF images. **(b)** The SAM-Plus GUI for GPU-based batch segmentation of representative worms, with results exportable as a zip file and interactive correction or manual segmentation for coiled worms. **(c)** VNC-Quant GUI, which maps normalized neuron positions along the VNC’s AP axis and generates a spreadsheet of neuron positions (DDs, DBs, DAs, SABs) per genotype, with Tabs offering various data visualizations.

All components of VNC-Dist were implemented in Python, with core functions built upon open-source libraries (SciPy [31],https://scipy.org/; NumPy [32],https://numpy.org/ for numerical analysis; scikit-image [33],https://scikit-image.org/; Pillow [34],https://python-pillow.org/; OpenCV [26],https://opencv.org/ for image processing; pandas [35],https://pandas.pydata.org/; Matplotlib [36],https://matplotlib.org/ for output visualization). To make the pipeline easier to use, we packaged VNC-Dist as a graphical user interface (GUI) using the Python-based Streamlit framework [37] (https://streamlit.io/), deployed within a Docker container on Linux/Ubuntu 24.04 LTS. This pipeline begins with a modified deep learning model, SAM-Plus, for segmenting 2D TIFF images of worms from noisy fluorescent micrographs. This model, which comprises approximately 636.12 million parameters, operates on a CUDA-enabled GPU on a compute cluster or on a dedicated GPU (e.g., GeForce RTX) to batch process images and accelerate throughput. The VNC-Dist pipeline includes two user-friendly GUIs (Fig. 3b):

1. SAM-Plus (GPU-based), an optimized version of SAM for precise worm segmentations, output as PNG masks.
2. VNC-Quant (CPU-based), a custom mathematical algorithm for mapping the neurons along the *VNCL1*, (generating annotated PNG images, and then calculating normalized relative distances as percentages, which it outputs as both individual and concatenated CSV files. It also provides different tabs for visualizing the data and for statistical analysis.

VNC-Quant and SAM-Plus each launch in a web browser at distinct ports (http://localhost:8501 for VNC-Quant and http://localhost:8502 for SAM-Plus), accessible via command line or desktop launcher. Both GUIs are fully compatible with macOS, Linux, and Windows through Docker, and support batch processing for rapid, and efficient high-throughput data analysis.

#### 2.2.2 Details of Worm segmentation with SAM-Plus

We integrated Segment Anything Model (SAM) [38] with the ViT-H image encoder into VNC-Dist without additional training (Fig. 4a). SAM provides two segmentation modes: automatic (grid-based) and interactive (user-prompted). To achieve full automation, we used automatic mode.

**Fig. 4.**
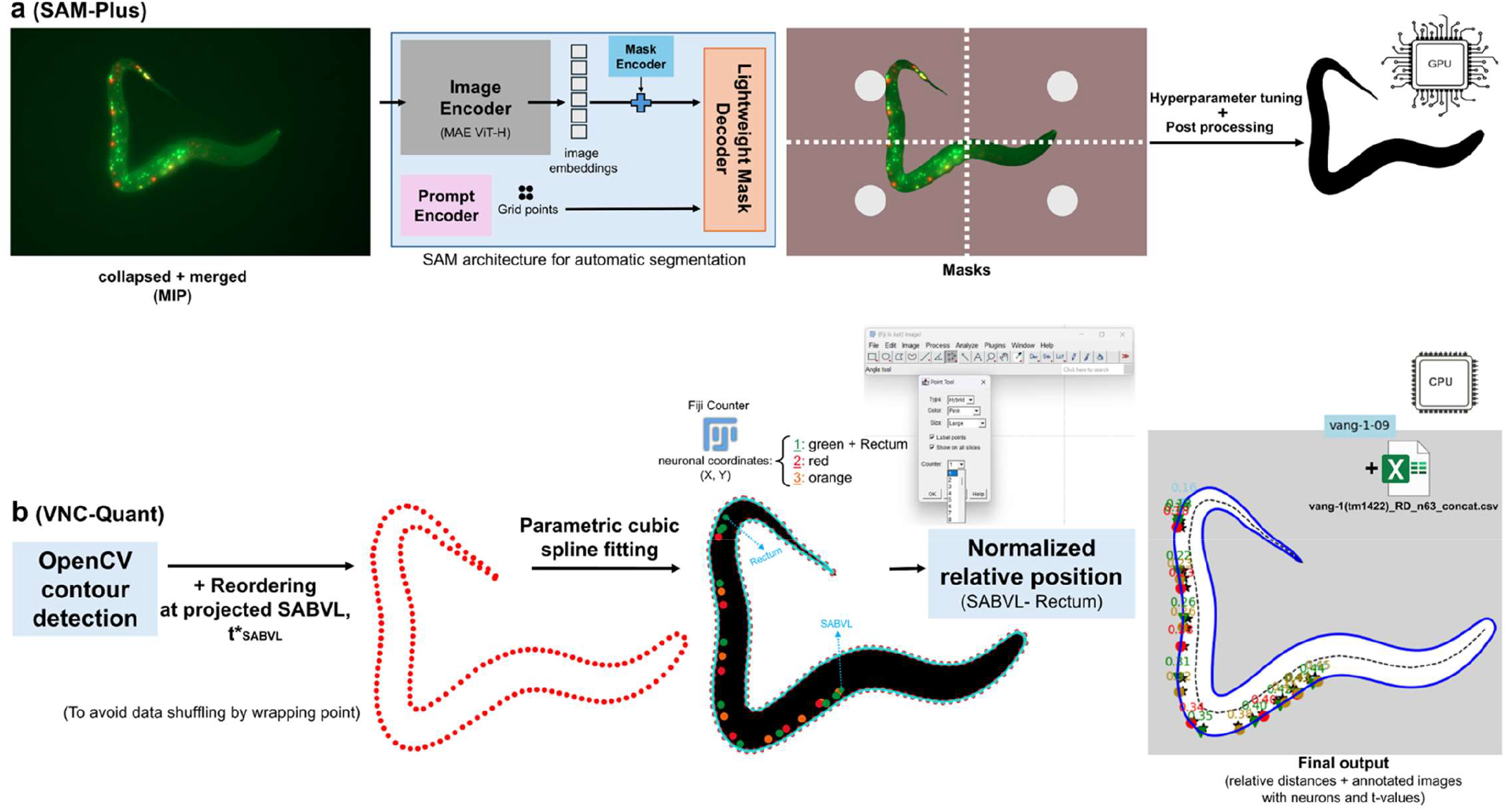
Overview of VNC-Dist pipeline for analyzing VNC neuronal positioning. CZI images are first compressed and merged, and neuron coordinates are extracted using Fiji’s multipoint tool (Counters: 1 for green points with rectum, 2 for red, 3 for orange). These data are then processed with VNC-Dist in two steps: **(a)** SAM-Plus automatically segments the worm body using the Segment Anything Model (SAM), prompted by a grid of two points per side, followed by post-processing to isolate and enhance the worm contour; and **(b)** VNC-Quant uses OpenCV to extract the pixel-level worm outline, fits a parametric cubic spline S(t) to these points, and projects each neuron onto this VNCL1 spline. Neuron positions are calculated via adaptive Gaussian quadrature, from the anterior (projected SABVL, 0%) to the posterior (rectum, 100%).

The automatic SAM mask generator was configured with two grid points per side to guide attention toward the worm, a prediction IoU threshold of 0.98, a stability score threshold of 0.98, one crop layer, a downscale factor of 1 (no downsampling), and a minimum mask region area of 99,000 pixels to discard smaller regions. We processed batches of 2D merged and collapsed micrographs as inputs for SAM. Initial SAM-generated masks were binarized and inverted to emphasize the foreground, then smoothed with a 21×21 Gaussian blur, a 33×33 median filter and a bilateral filter (d=9, σColor=75, σSpace=75), and finally subjected to morphological erosion (19×19 kernel) followed by closing (21×21 kernel). The refined segmented worms were saved as PNG files named to match their corresponding images. Due to the computational intensity of the process, an independent GUI version of the segmentation model, “SAM-Plus,” was developed using a Docker container and the Streamlit Python package (Fig. 3b). This version was specifically optimized for execution on a dedicated GPU. Although imaging coiled worms is not recommended, separate tabs in the SAM-Plus GUI allow for interactive correction of the segmented worms.

#### 2.2.3 Contour detection and spline fitting with VNC-Quant

VNC-Quant constructs a spline S(t), a smooth closed curve in image coordinates, that represents the boundary of the worm. First, OpenCV’s contour finder is applied to the SAM-Plus binary mask, producing a counterclockwise list of *N* control points

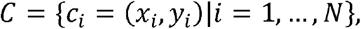

which by default begin at the top-leftmost pixel (Fig. 4b). To ensure the spline starts at the SABVL landmark, the boundary points *C* are circularly shifted before fitting any splines. The pipeline computes:

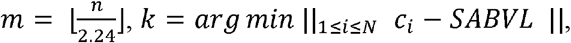

where the divisor 2.24 is an empirically estimated ratio of L1 VNC length to full worm perimeter. Thus, the pipeline replaces *C* with the sequence

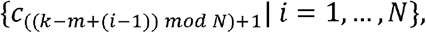

shifting the list by *k − m* positions so that the first point lies at SABVL. This guarantees motoneuron projections follow a continuous anterior-to-posterior order without wrap-around. Next, each (shifted) point is assigned the uniformly spaced parameter

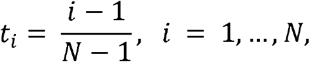

and fitted separate cubic splines *x*(*t*) and *y*(*t*) through {(*t*_*i*_,*x*_*i*_) and {(*t*_*i*_,*y*_*i*_), respectively, using SciPy’s FITPACK solver with a curvature penalty. The resulting smooth closed curve

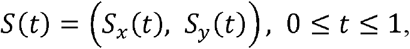

provides a reproducible boundary trace, starting at SABVL, for projecting neuronal coordinates and computing normalized distances along the VNC.

The fitted spline below is defined by minimizing the sum of squared deviations from the *N* contour points plus a λ-weighted curvature penalty and is computed in *O(N)* time using SciPy’s FITPACK cubic smoothing spline solver. Here, λ= 900 was chosen by visual inspection to balance fidelity to the sampled control points against penalization of curvature, producing a smooth boundary that is not required to interpolate every *c*_*i*_.

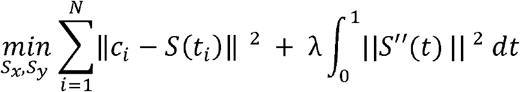

Let MN = (MN_x_, MN_y_) be the image coordinates of a given motoneuron. We projected it onto the spline S(t) by finding

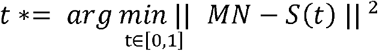

so that its spline coordinates are

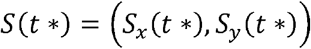

In particular, S(t*_SABVL_) locates the foot of the perpendicular from the SABVL landmark onto S(t). Because all motoneurons lie on the worm’s ventral surface, we defined the *VNCL1* spline as the segment of the closed curve S(t) that lies closest to their projected coordinates. To measure the distance along this ventral spline from SABVL at (at t*_SABVL_) to any other neuron at (at t*_*k*_), we numerically integrate the arc length

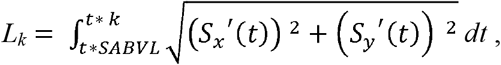

using SciPy’s adaptive Gaussian quadrature with an absolute tolerance of 10^−6^ and a maximum of 100 subintervals (Fig. 4b).

Therefore, if we have a sequence of *K* motoneurons whose projections onto the spline occur at

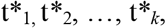

we first set

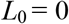

at the SABVL reference point. Then the arc length between the (*i* - 1)th and *i*th motoneuron is

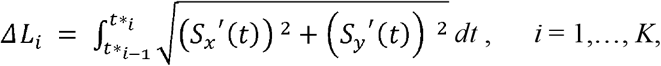

and the cumulative distance from SABVL up to the *K*th motoneuron is

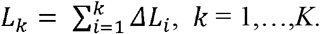

Finally, the normalized percent□of□path for the *k*th motoneuron is

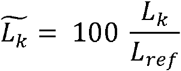

where *L*_*ref*_ is the total ventral□spline length from SABVL to rectum.

### 2.3 Statistical analysis

All analyses and graphing were performed in Python 3.10 using custom in-house scripts, which are also available through the user-friendly VNC-Dist GUI. Normality of neuron position distributions in wild-type (WT) and mutant strains was assessed via a three-step approach, box– violin plots, Q–Q plots, and the Shapiro–Wilk test (p ≥ 0.05). In the Q–Q plots, all genotypes were overlaid per neuron and compared by their fitted lines *y*= (*slope*)*x*+ (*intercept*), where the intercept approximates the neuron position mean and the slope the standard deviation (Figs S1–S3).

Pairwise WT versus mutant comparisons (n > 30 per group) employed two □sample t-tests: Student’s t-test when Levene’s test indicated equal variances, or Welch’s t-test otherwise. Effect sizes were quantified by Cohen’s d, with d ≥ 0.7 deemed significant. This effect size threshold corresponds to the minimal positional variability observed for DA8 and DA9 in PCP and *sax-3*/*Robo* mutants. For WT neurons, we built a correlation network using Pearson’s r ≥ 0.7 (p ≤ 0.05) to identify spatial clusters of motoneurons. To control for multiple comparisons in the VNC extension defect analyses, p-values were adjusted using the Bonferroni correction.

Finally, to evaluate performance across user experience levels, we computed the three-way mean absolute deviation (MAD) of neuron position measurements and processing time to assess inter-user variability, and both Pearson and intraclass correlation coefficients (ICC) to measure consistency between manual and VNC-Dist measurements. The overall mean absolute error (MAE) was also calculated to quantify accuracy.

## 3. Results

### 3.1 Multiplexed fluorescence imaging of DA, DD, and DB motoneurons

To label the DD, DA, and DB motoneuron classes at the L1 stage, we used transcription factors with class-specific expression: *unc-30* in DD1–6, *unc-4* in SAB and DA1–9, and *vab-7* in DB1–7 neurons [39–41]. Promoter sequences from these genes were used to drive red and green fluorophores, enabling distinct nuclear labeling of each motoneuron class. We used the MosSCI procedure [insert reference 37] to insert a single copy of an *unc-30* promoter-driven *mCherry::H2B* transgene into the *ttTi5605* site on chromosome II. CRISPR/Cas9 gene editing was used to endogenously tag the C-terminus of *unc-4* with mNeonGreen (mNG) and the C-terminus of *vab-7* with a dual-colour tag, *mNG::T2A::mScarlet-I::H2B*. The T2A self-cleaving peptide [42] enables co-expression of mNG and mScarlet markers. Histone H2B fused to mScarlet-I and mCherry was used to localize these fluorescent proteins to nuclei. In addition to DAs, *unc-4* is also expressed in three SAB neurons, SABVL, SABVR, and SABD, located in the RVG [41]. When used in WT and in *vang-1, prkl-1, sax-3*, and double mutant combinations, this reporter system enables visualization of DD nuclei in red, SAB and DA nuclei in green, and DB nuclei in yellow at the L1 stage (Fig. 1a; Fig. 6a).

### 3.2 Overview of the VNC-Dist pipeline

Figures 1–3 establish the foundation for our approach: Figure 1 illustrates the anatomical landmarks and spline model for motoneuron mapping, Figure 2 compares manual Fiji measurements with our semi-automated tool, and Figure 3 details the VNC-Dist GUI. Building on this groundwork, VNC-Dist offers a robust and standardized pipeline for the quantification of VNC neuronal positioning. This pipeline comprises two components. First, GPU-accelerated batch processing segments worms in micrographs using the state-of-the-art Segment Anything Model (SAM) (Fig. 4a). Second, CPU-based processing measures the normalized relative position of projected DD, DA, and DB nuclei along the *VNCL1* spline, which is on the ventral side of S(t), using SABVL (anterior) and the rectum (posterior) as reference points (Fig. 4b). Establishing these consistent landmarks enables reliable genotype comparisons, as normalization accounts for size variability among L1 worms.

To validate the relative distance measurements, the toolkit produces annotated images that display the projection of neurons onto the ventral side of the worm (*VNCL1*) along with the corresponding t-values for each neuron. This visualization confirms whether the ordering of t-values accurately reflects the anterior-to-posterior sequence of the projected neurons according to the reference points (Fig. 4b). In addition, VNC-Dist supports batch processing. The pipeline process is as follows:

First, neuronal coordinates are determined using the Fiji multipoint tool by manually selecting the centers of nuclei in 3D z-slices. Three distinct “Counter” numbers are assigned per class (1: DAs, SABs, and rectum; 2: DDs; 3: DBs), and the extracted X-Y coordinates are saved as CSV files (Fig. 4b). Next, worm images are segmented using the SAM-Plus. The VNC-Quant, a mathematical algorithm, then extracts the worm’s contours and fits a smooth cubic spline, S(t). Neurons are projected onto the spline segment between the SABVL and rectum, termed *VNCL1*, and their positions are normalized as percentages of the curve length (with SABVL at 0% and the rectum at 100%). This method accurately reflects the actual ventral nerve cord, which runs parallel to *VNCL1* (Fig. 1b).

### 3.3 Performance evaluation

To evaluate the efficacy of worm segmentation in the VNC-Dist pipeline, we applied it to images of WT, *vang-1, prkl-1*, and *sax-3* single and double-mutant. The pipeline successfully segmented over 290 images using SAM-Plus, and VNC-Quant quantified spline-assigned positions for more than 7,000 DD, DA, and DB nuclei relative to SABVL and rectum. Before conducting further analysis of VNC neuronal positioning to investigate VNC assembly, we manually reviewed each annotated image to ensure correct neuron projection and accurate anterior-to-posterior ordering of t-values. This manual review confirmed that all neuron projections and t-value orderings were correct and accurate, ensuring the validity of our subsequent analyses.

We next compared manual neuron position measurements with those obtained with VNC-Dist. To compare processing-time, we performed measurements of *prkl-1(ok3182)*, a mutant with strong neuron position defects [10]. This analysis showed that measurements obtained with VNC-Dist were on average 1.78 times faster than the manual method across beginner, intermediate, and expert users. Inter-user variability in processing time, quantified by three-way mean absolute deviation (MAD), decreased from 25.3 minutes with manual analysis to 7.1 minutes with VNC-Dist (a 3.6-fold reduction). Likewise, three-way MAD in neuron-position measurements across users was just 0.27 % with VNC-Dist, compared with 0.42 % for the manual method, indicating substantially greater inter-user variability in manual measurements. Three-way Pearson correlation and intraclass correlation coefficient (ICC) values averaged 0.99, reflecting excellent agreement between the neuron position measurements of the two methods. The overall mean absolute error (MAE) between manual and VNC-Dist neuron position measurements averaged 1.1%.

Together, these metrics demonstrate the speed, precision, and reproducibility of our semi-automated tool for high-throughput analysis.

### 3.4 Key outputs from VNC-Dist

We used VNC-Dist to quantify motoneuron positions in the WT VNC at L1. As expected, the neuronal positions in WT exhibited normal distributions, as confirmed by the Shapiro-Wilk test (p ≥ 0.05) and further supported by the Q–Q plot and violin-box plot analyses (Figs. S1–S3). This data allowed us to quantify multiple aspects of motoneuron positioning and patterning in WT. First, plotting the mean positions of DD, DA, and DB nuclei relative to SABVL and the rectum confirmed the consistency in their positioning along the VNC (Fig. 5a, b). Notably, DA and DB cell bodies (nuclei) are more widely spaced in the middle of the VNC and closer to each other at either end, whereas the DD cell bodies are roughly equally spaced regardless of where they occur in the VNC (Fig. 5c). Second, plotting the sequential order of motoneurons from anterior to posterior confirmed that their arrangement along the AP axis is highly stereotyped. Despite some variability in the most anterior DD and DB positions and excluding the most posterior DA8 and DA9, which lie opposite each other on the left and right, the DD, DA, and DB cell bodies display a consistent alternating pattern, with no two motoneuron classes positioned next to each other (Fig. 5d). Correlation network analysis (P < 0.05, Pearson r ≥ 0.75), which quantifies pairwise spatial similarity, delineated three clusters matching the VNC’s anatomical subdivisions. The VNC neurons formed one cluster with minimal variation in the central region of the network, while the remaining two clusters consisted of neurons in the RVG and PAG (Fig. 5e). These observations suggest that the positioning of middle VNC neurons may involve mechanisms distinct from those regulating neurons in the more tightly packed RVG and PAG regions.

**Fig. 5.**
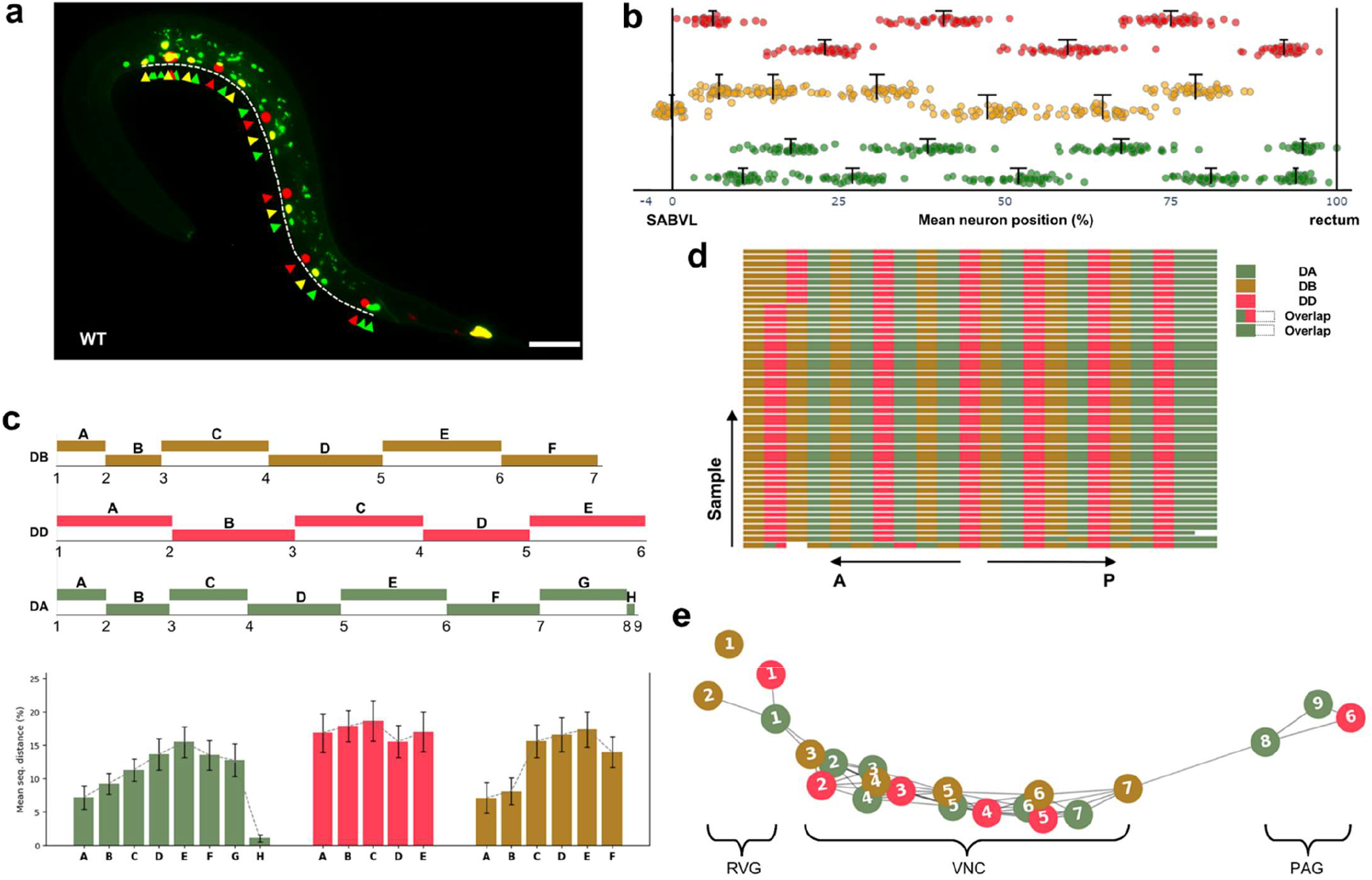
VNC Neuronal positioning in wild-type *C. elegans* analyzed by VNC-Dist. **(a)** Merged and collapsed micrograph showing motoneurons along the AP axis (dashed line). Green arrows: DAs and SABs; red arrows: DDs; orange arrows: DBs. Scale bar, 15 µm. **(b)** Dot plot of mean neuron positions normalized to SABVL (0%) and rectum (100%) for n = 42. Error bars represent mean ± 95% CI. **(c)** Mean sequential spacing of neurons along the AP axis, with segmented bar plots (top). Error bars represent SD. **(d)** Diagram showing the anterior-to-posterior order of all 22 motoneurons, grouped by class. **(e)** Correlation network analysis using motoneurons as nodes and significant relative distances (p < 0.05, Pearson r ≥ 0.75) as edges to group neurons into VNC, RVG, and PAG relative to SABVL (0%).

We next extended our analysis to PCP and Robo pathway mutants, which are known to affect motoneuron positioning in the VNC [10]. To assess the impact of *vang-1, prkl-1*, and *sax-3* mutations on neuronal positioning, we used our semi-automated pipeline to quantify the mean position (with 95% confidence intervals) of neurons relative to SABVL and the rectum in the following strains: *vang-1(tm1422), prkl-1(ok3182)*, and *sax-3(zy5)* single mutants, as well as *vang-1(tm1422) sax-3(zy5)*, and *vang-1(tm1422); prkl-1(ok3182)* double mutants. Violin–box plots and Q–Q plots were used to examine neuronal position distributions across mutant strains, with some approximating a normal distribution and others showing departures from normality (Fig. S1; Fig. S2). DA neurons, and to a slightly lesser extent, DB neurons, may remain normally distributed, signaling robust guidance, while DD neurons often deviate, revealing greater sensitivity to the PCP and Robo pathways (Fig. S2).

It is important to note that motoneurons of each class are numbered sequentially from anterior to posterior based on their plotted position and not by terminal lineage identity, which may be unknown in mutants due to position shifts. As expected, compared to WT, PCP and Robo mutants exhibited pronounced disruptions in neuronal positioning and organization (Fig. 6a, b). *vang-1* and *prkl-1* single mutants exhibited significant anterior shifts in the mean positions of many DD, DA, and DB cell bodies, with *prkl-1* mutants showing significantly more severe defects than *vang-1* mutants. These shifts were less pronounced in *sax-3* single mutants, with only DD2 and DD6 showing significant anterior shifts in their mean positions (Fig. 6b). However, an examination of individual neuron position distributions (Fig. S1) reveals anterior displacements in a subset of neurons, such as DD4, that are not evident when considering mean position values. In all three mutant backgrounds, these positional defects are accompanied by changes in the arrangement or pattern of motoneurons within the VNC of individual worms (Fig. 6c). In *vang-1; prkl-1* double mutants, the anterior shifts in mean neuron position more closely resemble the milder phenotype observed in *vang-1* single mutants than the stronger defects seen in *prkl-1* mutants, suggesting that, for some aspects of VNC assembly, VANG-1 acts downstream of PRKL-1. This epistatic interaction was noted previously when only DD neuron positions were assessed [10].

**Fig. 6.**
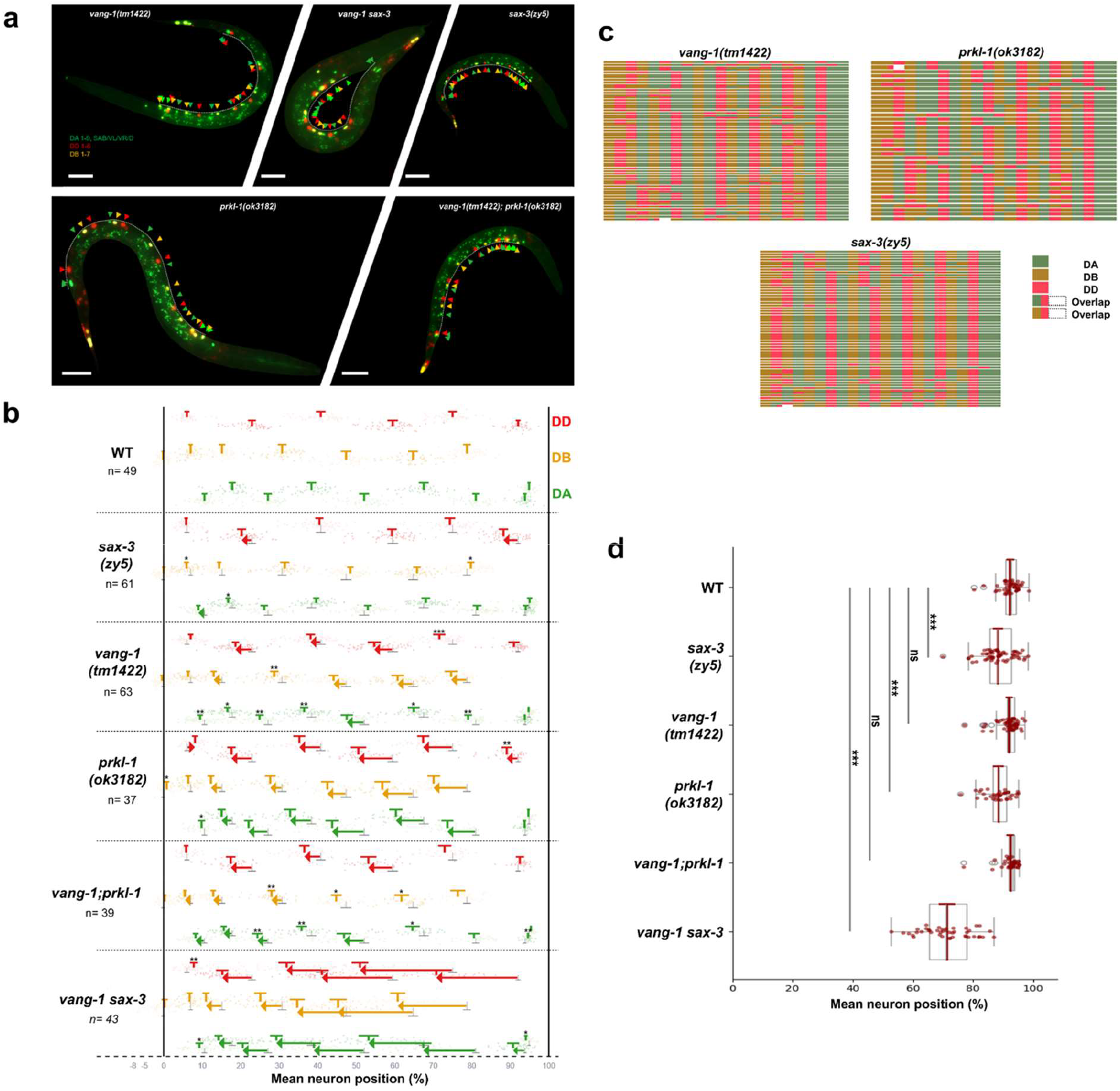
Quantification and analysis of VNC neuron displacements in single and double mutants via VNC-Dist. Representative fluorescent images of *vang-1(tm1422), vang-1(tm1422) sax-3(zy5), sax-3(zy5), prkl-1(ok3182)* and *vang-1(tm1422); prkl-1(ok3182)* mutants respectively. Green arrows: DAs and SABs; red arrows: DDs; orange arrows: DBs. The dashed white line represents the VNCL1 curve. Note: Worms were segmented from the background; Scale bar, 15 µm. **(b)** Dot plot of mean neuron positions normalized to SABVL (0%) and rectum (100%). Mean position of DD (red), DA (green), and DB (orange) neurons are compared to WT. Arrows indicate statistically significant, high-magnitude shifts (Cohen’s d ≥ 0.7) from WT to mutant. Significance is indicated by asterisks (* and **) and arrows (***). Error bars represent mean neuron positions ± 95% confidence intervals (CI); ***P < 0.001, **P < 0.01, *P < 0.05. **(c)** Diagrams showing the anterior-to-posterior order of all 22 motoneurons in single mutants, grouped by class. White spaces indicate overlapping neuron positions. **(d)** A measure of VNC extension is plotted as the mean position of the last neuron (excluding DA8 and DA9) relative to SABVL (0%) and rectum (100%) for each genotype (which, in these PCP and *sax-3*/*Robo* mutants corresponds to DD6). Pairwise comparisons between each mutant and WT were performed using Welch’s t-test with Bonferroni correction (α = 0.05). Significance levels are indicated as ***p < 0.001; ns, not significant. Mean ± 95% CI shown as red T shape. Red T-bars indicate mean ± 95% CI. Grey T-bars in mutants indicate WT mean ± 95% CI.

As expected, disruption of both the PCP-like and Robo pathways in the *vang-1; sax-3* double mutants showed the most pronounced anterior shifts, consistent with severe convergent extension defects during embryonic VNC assembly that likely interfered with the posterior migration of neuronal progenitors. This convergent extension defect is evident in a plot of the mean position of the most posterior motoneuron, excluding DA8 and DA9, which appear less affected by the disruption (Fig. 6d).

These results indicate that VNC-Dist can replicate previous findings from manual measurements, which focused exclusively on DD neuron positions, but in a way that is less labor-intensive and likely more reproducible across users. Furthermore, by examining all L1 VNC motoneurons, it extends these observations to include DA and DB neurons, providing a more comprehensive view of the perturbations occurring during embryonic VNC assembly.

## 4. Discussion

Taken together, our microscopy and software pipeline for quantifying neuron cell body positions within the L1 VNC provides precise reproducible measurements crucial for understanding genetic factors influencing neuronal organization and positioning in C. elegans. This high-throughput interactive pipeline integrates deep learning with numerical analysis methods to robustly quantify neuron spacing along the VNC.

Our pipeline analyzed fluorescence micrographs from various mutant strains, including *vang-1, prkl-1, sax-3*, and their double mutant combinations, revealing distinct patterns of neuronal displacement. These findings underscore the critical role of these genes in establishing the spatial order of motoneurons during VNC assembly. VNC-Dist is a niche pipeline that is distinct from other generalist platforms such as NeuroPAL [43] or CellProfiler [44], because it is optimized for VNC motoneuron positioning at L1 stage. However, VNC-Dist can be adapted to analyze cell positioning in *C. elegans* at other developmental stages, quantifying ventral, dorsal, and midline cell positioning along the AP axis. By using parametric spline fitting with numerical analysis for continuous curve length calculation, and deep learning for worm outline extraction, the method yields robust and precise quantification across diverse worm shape and cell positioning scenarios. The VNC neurons were projected onto the ventral outline rather than the spine line [45], as it offers both true absolute cell-to-cell distances and normalized distances based on the anterior and posterior reference points (SABVL as 0% and rectum as 100%), providing a more biologically representative model of the VNC.

The primary objective of this study was to develop VNC-Dist, a user-friendly tool designed to replace the time-consuming, laborious, and error-prone manual measurement of VNC motoneuron positions. We envision further enhancement of the VNC-Dist into a fully automated pipeline. Using both state-of-the-art nuclei segmentation [46] and classical image processing techniques, future versions will aim to identify automatically all motoneuron nuclei at scale, including DA, DD, and DB types, thereby eliminating the need for manual annotation in Fiji to extract the positional information. Furthermore, the segmentation component of VNC-Dist, SAM-Plus, does not require training dataset generation or model training. However, it can be fine-tuned using the mask decoder of Meta AI’s SAM algorithm with human-verified ground truth generated by SAM to facilitate the segmentation of challenging coiled worms.

Consequently, VNC-Dist offers a powerful platform that combines mathematical precision with deep learning to produce accurate results in a time-efficient manner. High-throughput analysis of VNC neuron types from diverse fluorescence images and genotypes advances our understanding of the molecular and cellular mechanisms that govern neuronal positioning.

## Code availability

The reviewed version of the VNC-Dist pipeline, including the code, GUI, and future updates, is available on GitHub under a permissive open-source license at https://github.com/C-elegans-VNC/VNC-Dist. Additionally, test data, including TIF images and the corresponding CSV coordinates, were provided to facilitate testing and evaluation of the pipeline.

## Acknowledgements

Some strains were provided by the Caenorhabditis Genetics Center, which is funded by the National Institutes of Health Office of Research Infrastructure Programs (P40 OD010440). The authors acknowledge the Cell Biology and Image Acquisition Core (RRID: SCR_021845) funded by the University of Ottawa, Ottawa, Natural Sciences and Engineering Research Council of Canada (NSERC), and the Canada Foundation for Innovation. This work was supported by grants from the Canadian Institutes of Health Research (CIHR FRN 123513 and 156160) and NSERC (2018-06790) to A. Colavita, and an NSERC grant (2019-06604) to T. J. Perkins.

