## Supplemental Figures for "VNC-Dist: A machine learning-based semi-automated pipeline for quantification of neuronal positioning in the *C. elegans* ventral nerve cord"

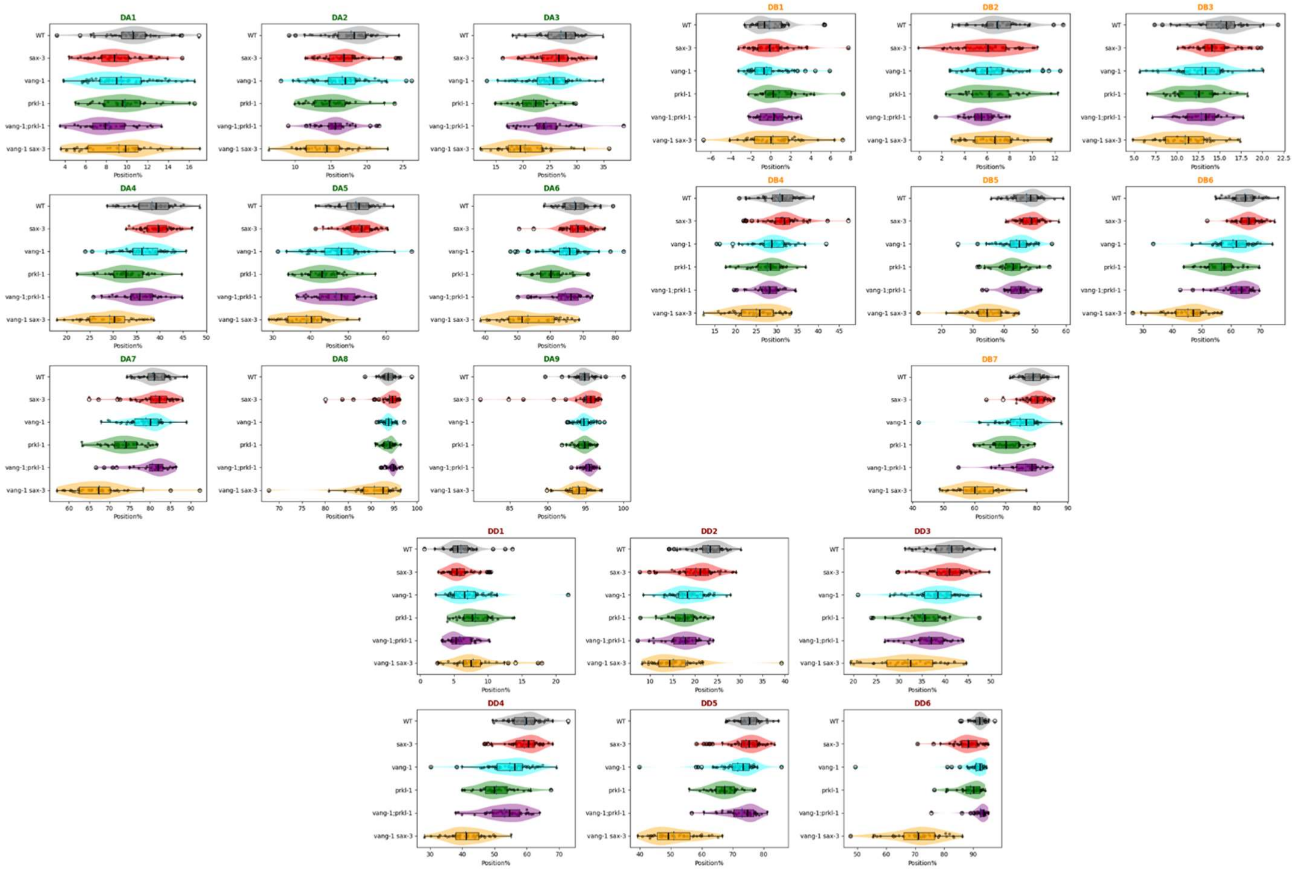


**Fig. S1 | Visualization (violin plot overlaid a box plot) of DD, DA and DB neuronal positions across genotypes**

Mutant strains (*prkl-1(ok3182)*, *sax-3(*zy5*)*, *vang-1(tm1422), vang-1;prkl-1, vang-1 sax-3*) exhibit altered anteroposterior neuron positioning compared to WT. Distributions assess normality and highlight genotype specific effects on neuron positioning along the AP axis, with median (black) and mean (blue) indicated by vertical lines.

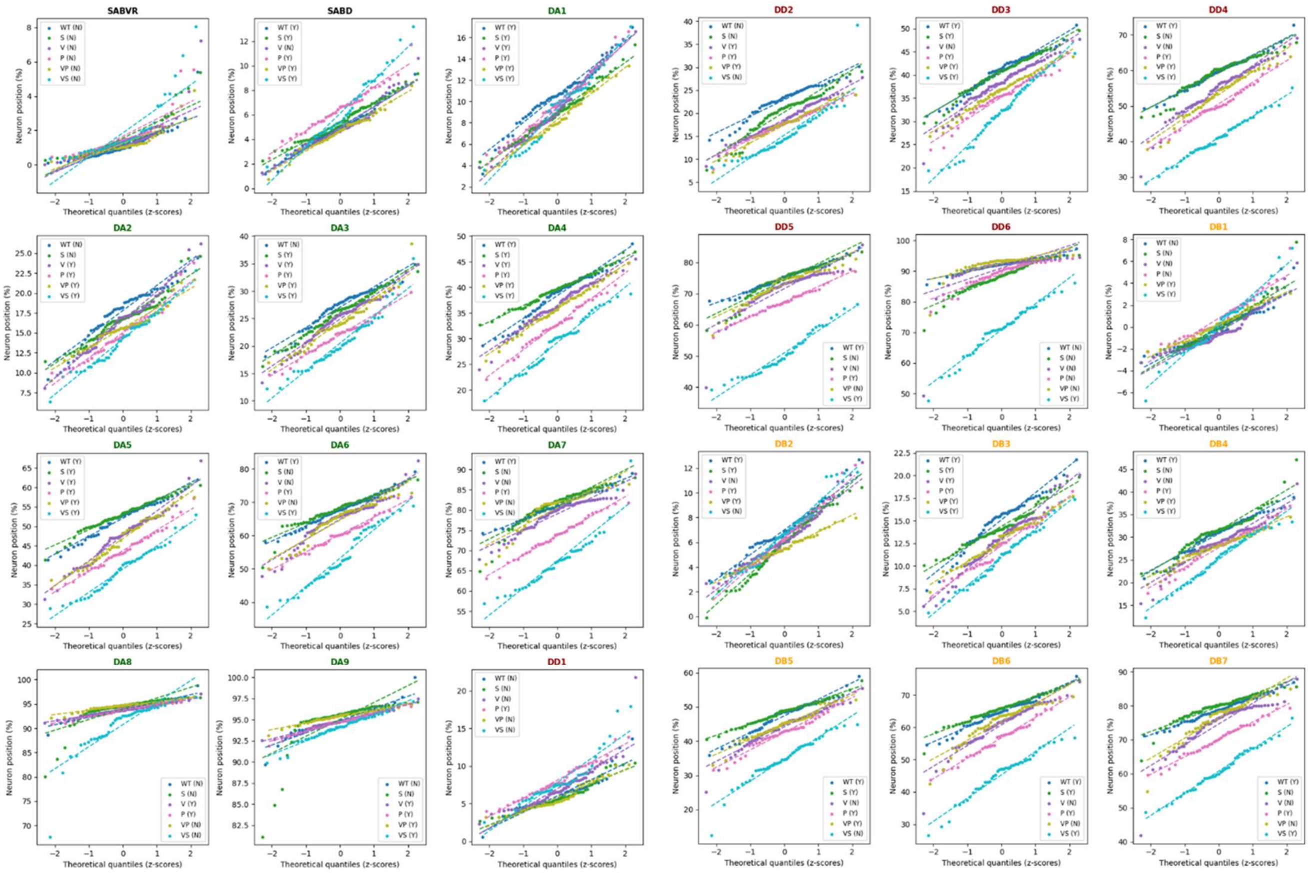


**Fig. S2 | Normality assessment of neuron positions for DA, DD, and DB neurons in WT and mutants**

Q–Q plots with Shapiro–Wilk tests assess the normality of DA, DD and DB neuron position distributions in WT and five mutant genotypes (P: *prkl-1(ok3182),* V: *vang-1(tm1422),* S: s*ax-3(zy5),* VP: *vang-1;prkl-1,* VS: *vang-1 sax-3*). Deviations from the diagonal indicate non-normality. Legend entries “Genotype (Y/N)” report Shapiro–Wilk test outcomes (Y = normal; N = non-normal) and are color-coded by genotype. VS (cyan) consistently has the lowest intercept (most anterior shift) across neurons, with P (pink) a close second, and VS’s markedly steeper slope reflects its larger variance.

**a**

| Neuron | Genotype | W | P_Value | Normal |
|---|---|---|---|---|
| DA1 | WT | 0.974967 | 0.376849 | Yes |
| DA2 | WT | 0.94196 | 0.017642 | No |
| DA3 | WT | 0.949236 | 0.034397 | No |
| DA4 | WT | 0.965363 | 0.157332 | Yes |
| DA5 | WT | 0.974481 | 0.361376 | Yes |
| DA6 | WT | 0.987908 | 0.891802 | Yes |
| DA7 | WT | 0.983884 | 0.73378 | Yes |
| DA8 | WT | 0.951555 | 0.042702 | No |
| DA9 | WT | 0.90887 | 0.001084 | No |
| DB1 | WT | 0.877562 | 0.00011 | No |
| DB2 | WT | 0.967022 | 0.183804 | Yes |
| DB3 | WT | 0.962407 | 0.119016 | Yes |
| DB4 | WT | 0.970267 | 0.248143 | Yes |
| DB5 | WT | 0.961642 | 0.110691 | Yes |
| DB6 | WT | 0.981502 | 0.629555 | Yes |
| DB7 | WT | 0.989455 | 0.936715 | Yes |
| DD1 | WT | 0.903033 | 0.000691 | No |
| DD2 | WT | 0.943885 | 0.021014 | No |
| DD3 | WT | 0.965686 | 0.162171 | Yes |
| DD4 | WT | 0.985925 | 0.819315 | Yes |
| DD5 | WT | 0.979769 | 0.555616 | Yes |
| DD6 | WT | 0.950895 | 0.040145 | No |

**b**

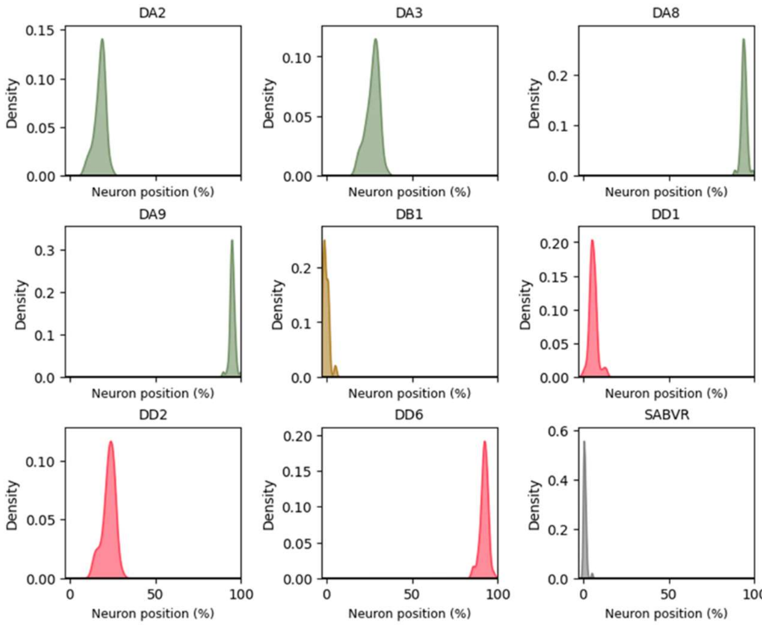


**Fig. S3 | Shapiro–Wilk test results for normality of neuron positions in WT**

**(a)** Neurons with p-values > 0.05 are considered normally distributed. **(b)** Although DA2, DA3, DA8, DA9, DB1, DD1, DD2, and DD6 showed significant deviations from normality ($p < 0.05$), their Shapiro statistics, histograms, and Q–Q plots suggest they can be reasonably assumed to follow a normal distribution (n= 49).
