## Supplemental Table 1 for "VNC-Dist: A machine learning-based semi-automated pipeline for quantification of neuronal positioning in the *C. elegans* ventral nerve cord"

### Supplementary information

**Supplemental Table S1. Primers used for CRISPR knock-ins.**

| | |
|---|---|
| ***unc-4(zy123[unc-4::mNG::3xFlag])*** | |
| guide | AGCGACTAATGCATTGACTAGTTTAAGAGCTATGCTGGAA |
| 5’unc4arm.F | acgttgtaaaacgacggccagtcgccggcaatttctccccgtcacgtcttc |
| 5’unc4arm.R | CATCGATGCTCCTGAGGCTCCCGATGCTCCTACACTTTTCAGTAATTCAGCAACAGTAGTCAATGCATTAGTCGCTAC |
| 3'unc4arm.F | CGTGATTACAAGGATGACGATGACAAGAGATAAatttttttaaaattcaattttgaaccgtgccc |
| 3'unc4arm.R | ggaaacagctatgaccatgttatcgatttcatttcagctctgcgagacgt |
| ***vab-7(zy137[vab-7::mNG::3xFlag])*** | |
| guide | TTAATCTGTAGAATAAGGCGGTTTAAGAGCTATGCTGGAA |
| 5'vab7arm.F | acgttgtaaaacgacggccagtcgccggcacaagtggccattactcgcaa |
| 5'vab7arm.R | CATCGATGCTCCTGAGGCTCCCGATGCTCCGTCGGTAGAATAAGGCGAGGGAGA |
| 3'vab7arm.F | CGTGATTACAAGGATGACGATGACAAGAGATAAttcactatttttggaaaaaaaatcgg |
| 3'vab7arm.R | ggaaacagctatgaccatgttatcgatttcgttgtggcaaattgataggaa |
| ***vab-7(zy142[vab-7::mNG::T2A::mScarlet-I::H2B])*** | |
| guide | GAGAATCTGTACTTTCAATCGTTTAAGAGCTATGCTGGAA |
| H2B_vab7c.F | ccaagtacacttccagcaagtgacgatgacaagagataattc |
| T2A_mNG.R | cctctgccctctccagatcccttgtagagctcgtccattc |
